# Molecular basis of the pH-controlled maturation of the tick-borne encephalitis flavivirus

**DOI:** 10.1101/2022.10.27.514041

**Authors:** Emmanuelle Bignon, Elise Dumont, Antonio Monari

## Abstract

Flaviviruses are enveloped viruses causing high public concerns. Their maturation spans several cellular compartments having different pH. Thus, complex control mechanisms are in place to avoid premature maturation. Here we report the dynamical behavior at neutral and acidic pH of the precursor of the membrane fusion protein E of tick-borne encephalitis, showing the different stabilization of the E dimer and the role played by the small fusion-assisting protomer (pr). The comprehension, at atomic resolution, of the fine regulation of viral maturation will be fundamental to the development of efficient strategies against emerging viral threats.

## Introduction

In the last years flaviviruses have emerged as a major public health concern.^[1,2]^ Such viruses, which can be hosted by different animal reservoirs^[3,4]^ and be transmitted to humans, are the causative agents of rapidly spreading pathologies of high severity and impact. These include, for instance, Zika,^[5]^ Dengue,^[6]^ West Nile,^[7,8]^ Yellow Fever,^[9]^ transmitted by mosquitoes’ bites, as well as the tick-borne encephalitis (TBE) virus.^[10]^ The latter belongs to the *Flaviviridae*, a family of enveloped positive-sense RNA viruses encompassing three genera.^[2]^ Interestingly, the members of the *Flavivirus* genus, including TBE, present a peculiar class II membrane fusion protein E,^[11,12]^ whose reversible and irreversible conformational evolutions are fundamental in assisting the maturation of the virus.^[13]^ Indeed, the E protein, which is a rather large β-barrel rich protein, harbors a specific loop which inserts into the cellular membrane to promote the fusion with its viral counterpart, thus allowing the insertion of the virus’ genetic material. Importantly, the steps of the viral cycle are taking place in different cellular compartments, which are characterized by different pH values.^[14,15]^ Indeed, the first budding of the viral particles takes place in the endoplasmic reticulum (ER)^[16]^ at neutral pH, and is followed by their maturation in the mildly acidic trans-Golgi network (TGN). After exiting to the extracellular medium at neutral pH, the virions may infect the target cell through the interaction with the cellular receptor, and further trigger the acid-mediated fusion of the viral and the endosomal membranes.^[17]^

Therefore, precise regulation mechanisms are essential to avoid the instability of the viral membrane due to premature fusion-triggering in the TGN.^[14,15]^ It has been shown that such mechanisms are driven by the drastic conformational transition of the membrane fusion protein E. The structure of the soluble domain of the E protein (sE) in different environments has been recently reported, contributing to the understanding of the molecular basis of the regulation mechanisms.^[18]^ Indeed, in its dimeric form sE prevents membrane fusion.^[19–21]^ Interestingly, the association of the sE dimer (sE_2_) with two smaller protomer units (pr) has been pointed out as the strategy developed by the virus to avoid premature fusion despite the low pH of the TGN.^[22–25]^

In particular at acidic pH, the tetrameric (pr/sE)_2_ form remains stable, while sE_2_ dissociates into the two constituent monomers.^[18]^ Analysis of the crystal structures have pinpointed the main interactions between pr and sE, which might drive the stabilization of the (pr/sE)_2_ tetramer. Furthermore, possible conformational transitions favoring the interactions between pr and sE in mildly acidic conditions (pH 5.5) have been highlighted. These involve the flexible 150 loop region of sE_2_ and might favor the exposure of a pr-binding site at low pH. Yet the precise molecular mechanisms behind the pH-dependent stabilization of the sE dimers are not yet fully understood, while controlling this process could lead to the development of novel antiviral strategies. In this contribution we resort to high-level, long-range molecular dynamic (MD) simulations exceeding the μs time-scale to unravel, at an atomistic level, the precise interactions network and conformational equilibrium of the sE_2_ dimer and the (pr/sE)_2_ tetramer as well as their pH dependency.

## Results and Discussion

Different initial model systems have been constructed involving the sE_2_ dimer and the (pr/sE)_2_ tetramer. MD simulations for the two systems have been performed either at a neutral pH of 7 or at an acidic pH of 5.5. The effects of the pH variation have been modelled by adapting the protonation state of the titratable residues, i.e. histidine and glutamic acid, also based on the pKa predictions from the propKa web server.^[26]^ Furthermore, the binding free energies between the two sE units have been estimated using the Molecular Mechanics Generalized Born Surface Area (MM/GBSA) method. Classical MD simulations have been performed in the isothermal and isobaric ensemble (NPT) at 300 K and 1 atm using NAMD software^[27,28]^ and analyzed with VMD.^[29]^ Protein residues were modelled using the Amber ff14SB force field,^[30]^ and a time step of 4 fs was used to propagate Newton’s equations of motion thanks to the combination of Hydrogen Mass Repartition (HMR)^[31]^ and Rattle and Shake algorithms. Additional computational details are provided in Supplementary Information (SI).

In Figure 1 we report the time series of the distance between the center of mass of the two sE units in the (pr/sE)_2_ complex, which remains remarkably constant along the MD trajectory, with fluctuations not exceeding 2 Å. Globally the same behavior can be observed at either neutral or acidic pH, with the complex interacting persistently. The same information conveyed by the analysis of the time series can also be appreciated pictorially by the representative snapshots, also reported in Figure 1, and showing the maintaining of the compact (pr/sE)_2_ structure.

**Figure 1.**
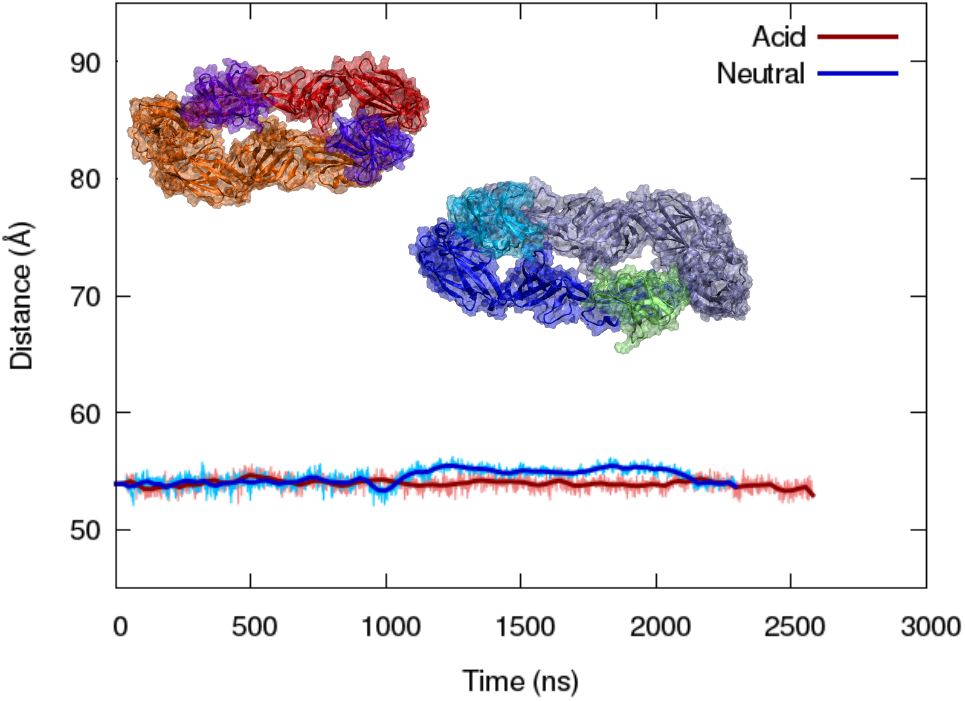
Time evolution of the distance between the centers of mass of sE units in the (pr/sE)_2_ at acidic (red) and neutral (blue) pH. Representative snapshots of (pr/sE)_2_ are also given in the inlay for acidic (left, red shades) and neutral (right, blue shades) conditions. pr are represented in violet and cyan for acid and neutral conditions, respectively, while the sE units are represented in red/orange and grey/blue.

The still rather limited time-scale of our MD simulations does not allow to observe the dissociation of the pr units even at neutral conditions. However, an in-depth analysis of our MD underlines the evolution of the interaction network as a function of the pH. As a matter of fact, the main residues participating to the stabilization of the tetramer have been identified by Vaney *et al*. from the corresponding crystal structure.^[18]^ Figure 2 provides a cartoon representation of the non-covalent, mainly salt bridge, interaction that enhance the stability of the (pr/sE)_2_ assembly, as obtained by our MD simulations. Note that because of the tetrameric organization, two symmetric interaction patterns between pr and sE can be evidenced. The full statistical analysis including whiskers plot defining the distribution of distances between the key amino acids bridging sE and pr are also given in SI.

**Figure 2.**
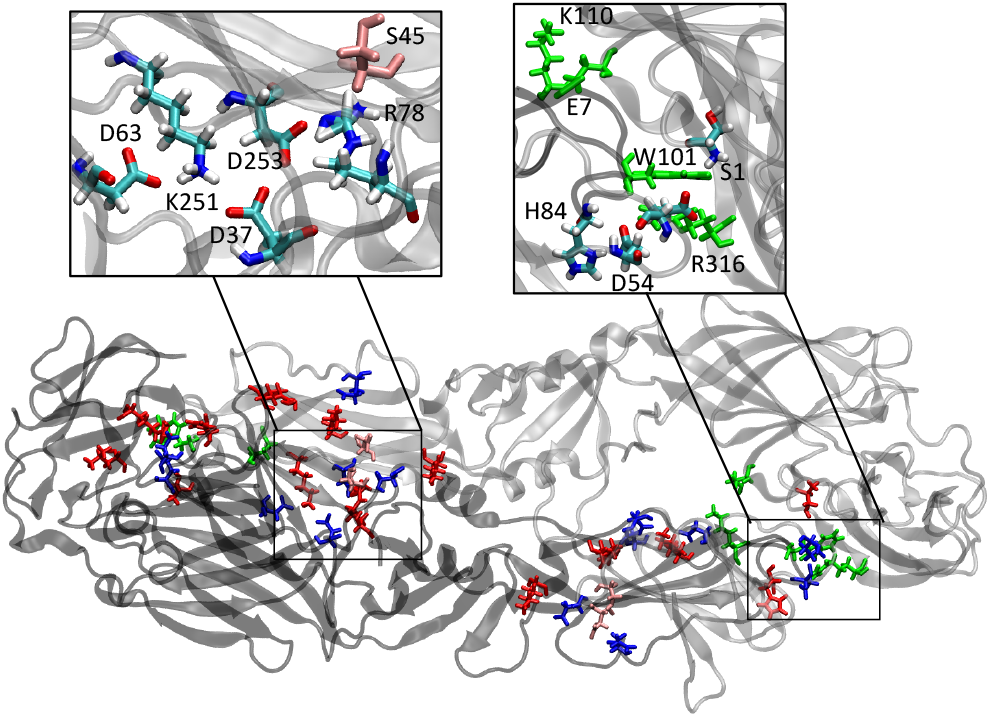
Cartoon representations of the (pr/sE)_2_ dimer, with residues implied in a network of cooperative salt bridges displayed in red and blue sticks respectively for positively- and negatively-charged residues. Residues shown in pink correspond to serine residues implied in an interaction. Residues in green correspond to sE/sE interactions. The inlays report zooms of some of the interactions.

With the exception of the salt bridges identified between E155(sE) and R68(pr) in the crystal structure, which are not preserved in our simulations, the global interaction pattern described by Vaney *et al*. is maintained. However, some interactions, such as K64(sE)/D43(pr) or D67(sE)/S45(pr) appear mobile and give rise to bimodal distributions oscillating between bound and unbound states. Figure 2 shows numerous positively- and negatively-charged residues at the pr/sE interface that corresponds to a cooperative network of seven pr/sE salt bridges, resulting in an enhanced stability compared to the sE_2_ homodimer. For the latter system, only two main non-covalent interactions come into play: a *π*-cation interaction between W101 and R316 and a salt bridge between E7 and K110, the latter being disrupted at acidic pH. Interestingly, the interactions involving either pr1/sE1 or pr2/sE2 correspond to six salt bridges, and are thus clearly stronger and more persistent than the cross terms between pr1/sE2 or pr2/sE1 (a sole salt bridge D57-R316 being identified with a distance lower than 4 Å).

Thus, although the two distributions are similar, a slightly stronger interaction network can be evidenced for the acidic medium, coherently with the experimentally observed stabilizing role of pr. Indeed, in the neutral environment we observe an important weakening of three interactions: K64(sE)-D43(Pr) with an increase by 2.5 Å, and H104(sE)-D54(pr), whose average distances is increased by ca. 4 Å. Furthermore, both the direct and cross-contact interactions are significantly reshuffled. As a matter of fact, we may also evidence that the increase in the distance distribution, and hence the weakening of the interaction network, is even more pronounced for the two cross contacts K64(sE)-D43(Pr) and H104(sE)-D54(pr), the histidine H104 interacting more strongly when double protonated. A deeper and aprioristic analysis of the interaction network based on the contacts developed along the MD simulations can be found in SI and confirms this analysis.

Globally, from the MD simulation of the (pr/sE)_2_ tetramer, we may conclude that the role of pr is fundamental in providing a number of pr-based key contacts bridging the two sE units, hence stabilizing the tetrameric structure. Furthermore, the interactions appear more favorable in an acidic environment rather than at neutral pH, suggesting that the formation of the tetramer could be controlled by the acidity level of the environment, hence could be dependent on the cellular compartment localization. The estimation of the binding free energies between the two sE monomers confirms this qualitative analysis: the presence of the pr units stabilizes the sE_2_ dimer at acidic pH but not at neutral pH (ΔΔG_binding_ values of −26 kcal/mol and 19 kcal/mol, respectively). More details of the free energy calculations and individual contributions of amino acids can be found in SI.

On the other hand, the MD simulations of the sE_2_ dimer, at acidic and neutral pH, reveal crucial and highly evident differences. Indeed, as shown in Figure 3, we see that in a neutral environment the sE_2_ dimer remains stable, exhibiting only a very modest evolution of the distance between the centers of mass of the two sE units. Conversely, in acidic medium a sharp increase of the distance between the two monomers is observed after 1.5 μs, which leads to the spontaneous and complete dissociation of the protein complex before the end of the MD simulation (2.5 μs). The striking different behavior in the two media is also illustrated by the final structures of the MD simulations (Figure 3B). A movie of the MD simulation in the acidic medium is provided in SI, allowing to appreciate the quick and irreversible disassembly of the sE_2_ dimer at low pH.

**Figure 3.**
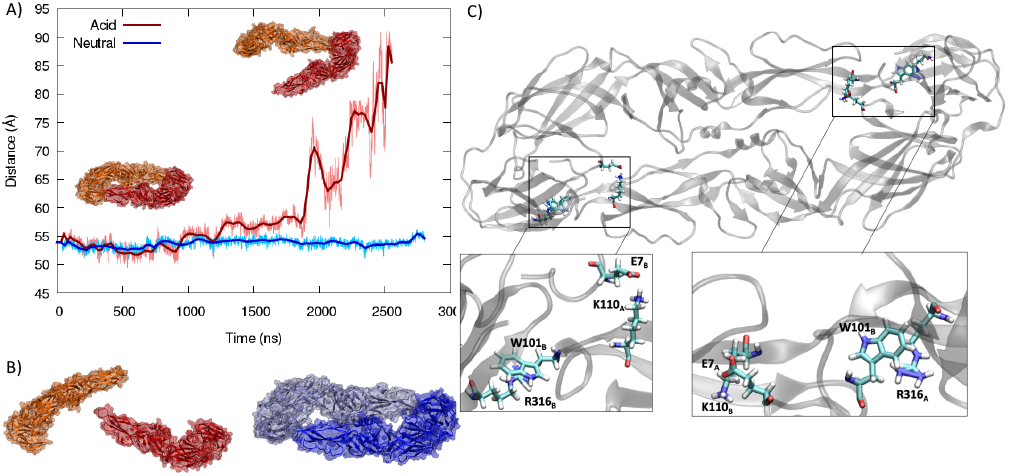
Time series of the centers of mass between the two sE units at acidic and neutral pH for the sE_2_ complex in the absence of pr (A). Representative snapshots showing the evolution of the complex in acid medium are given in the inlay. Final snapshot of the MD simulation for acidic (top, red colours) and neutral (bottom, blue colours) showing the stability of sE_2_ at neutral pH and the dissociation in acidic medium (B). Cartoon representations of the sE_2_ dimer, highlighting non-covalent interactions, i.e. E7/K110 and W101/R316, at the sE/sE interface (C).

These results, somehow striking considering the inertia due to the high mass of the protein complex, are coherent with experimental evidences obtained from high performance liquid chromatography^[18]^, in which the monomeric sE is eluted in acidic medium, while the dimeric sE_2_ form is observed at neutral pH. Coherently with the results presented in Figures 1 and 2, the tetramer (pr/sE)_2_ is instead stable and eluted even at low pH.

In addition to rather unspecific hydrophobic interactions developed at the sE/sE interface, only two symmetric, non-covalent interactions, namely one salt bridge E7/K110 and a *π*-cation interaction W101/R316, shown in Figure 3C, are present to consolidate the homodimeric sE/sE protein-protein interface. At neutral pH, these interactions maintain the two sE units yet the protonation of E7 is critical in triggering the unfolding of the sE_2_ dimer in acidic media. This is also highlighted by the destabilizing contribution of E7 and K110 to the sE_2_ ΔΔG_binding_ in these conditions, while W101 and R316 show stabilizing contributions – see SI. While the crucial difference in the stability of sE_2_ depending on the pH and the important role of pr in stabilizing the tetramer at low pH appear unambiguous, another crucial aspect remains to be pointed out. Indeed, it has been shown that the 150-loop of sE plays a fundamental role in fostering the interaction between sE and pr.

In Figure 4 we report representative snapshots and the evolution of the distance between the center of mass of the 150-loop and the rest of the corresponding sE unit at different pH. It is evident that a different dynamic is engaged in acidic or neutral conditions. Indeed, the average distance between the loop and the rest of sE in acidic conditions is clearly larger compared to the neutral medium. Remarkably, this behavior is confirmed for both monomers and remains evident even after the dissociation of the sE_2_ complex in acidic medium (Figure 4B and C). Interestingly, the evolution of the distance of the 150-loop from the rest of the sE monomer at low pH is closer to the one observed in the (pr/sE)_2_ complex (values in SI). This is in line with the exposure of the pr-binding area driven by the 150-loop at low pH, that might favor pr binding in acidic conditions. On the contrary, in neutral environment the loop is staying closer to the rest of the protomer, and most notably is forming a lid on the pr-binding surface. As a consequence, the 150 loop is, at neutral pH, partially shielding the pr interaction area (Figure 4A), hence adding a kinetic blockage to the formation of the (pr/sE)_2_ tetramer, which could also add up to the slight destabilization of sE_2_ binding energy in the presence of pr. Interestingly, this aspect was already pointed out experimentally, albeit considering only a static picture from the crystal state. Furthermore, in the (pr/sE)_2_ tetramer, the doubly protonated H146 (at acidic conditions) induces an overall translation of the loop, disrupting the interaction channels with S1(sE2) and E58(pr1). Our simulations coherently preview that the acidic environment favors the interaction between pr and sE_2_ in a two-fold fashion: thermodynamically, i.e. increasing the (pr/sE)_2_ binding affinity, and kinetically, facilitating the exposure of the pr interaction region.

**Figure 4.**
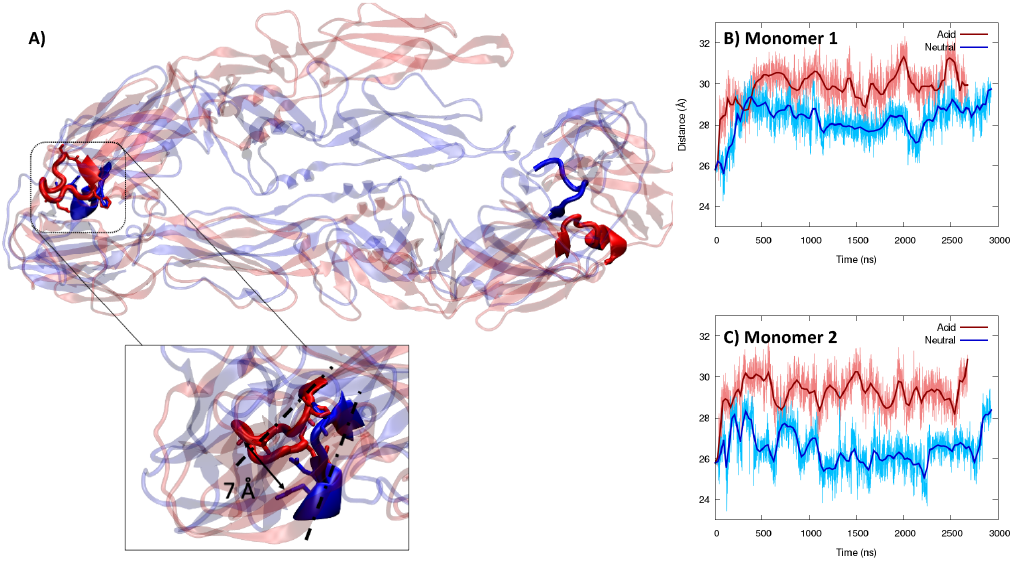
Superposition of the structure of the sE_2_ dimer at acidic (red) and neutral (blue) pH. The 150-loop is evidence using a solid representation and also highlighted in the inlay (A). Note that a snapshot showing a partial destabilization of the dimer has been chosen for the acidic pH. Time evolution of the distance between the 150-loop and the rest of the corresponding sE unit for the first (B) and the second (C) monomers, in the absence of pr.

## Conclusion

In the present contribution we have studied, by using all atom MD simulations exceeding the μs time-scale, the intrinsic dynamic of the soluble E (sE) protein of the TBE virus. Indeed, the maturation of the virion requires a precise regulation and adaption to variable pH conditions, due to the coexistence of differential maturation phases taking place in different cellular compartments.

In particular, we have shown that the interaction between sE and the pr protomer is, indeed, able to stabilize the formation of the complex at low pH. A more pronounced and persistent interaction network is observed between sE and pr in acidic conditions rather than at neutral pH. The stabilization of the (pr/sE)_2_ tetramer by pr in acidic environment is also confirmed by the estimation of the respective binding free energies. Strikingly, instead, the sE_2_ dimer is not stable at low pH, and a spontaneous dissociation is observed after 2 μs of MD simulation. These results confirm the finely-tuned regulation of the maturation cycle of TBE: the formation of the (pr/sE)_2_ tetramer is favored and maintained in acidic conditions by the pr unit, which develops favorable bridging interactions between the two sE units. In turn, the pr-driven stabilization of the complex avoids the premature insertion of the soluble E protein in the TGN apparatus, which would hamper the viral maturation. Interestingly, we have also shown that in acidic conditions the sE 150-loop is pushed further from the pr interaction region, hence removing kinetic constraints to the formation of the (pr/sE)_2_ tetramer. Conversely, in neutral conditions, where the sE_2_ dimer is inherently stable, the pr binding region is partially shielded by the 150-loop, reducing the probability of the formation of the complex.

Flaviviruses represent public concerns viral vectors, and different epidemic outbreaks are observed, also favored by global warming which further extend the regions accessible by viral vectors, mainly mosquitoes and ticks.^[32]^ Therefore, the fine comprehension of the replication cycle and of the different checkpoints that are established between different cellular compartments is fundamental. This approach is indeed also instrumental in assuring a rational design of putative antiviral agents capable of targeting different steps of the viral replication cycle. Altering the pr-based control of sE aggregation could result in an original therapeutic strategy against an important family of emerging pathogens.

## Acknowledgements

The authors thank GENCI and Explor computing centers for computational resources. E.B. thanks the CNRS and French Ministry of Higher Education and Research (MESR) for her postdoc fellowship under the GAVO program. The Platform P3MB is gratefully acknowledged. A.M. acknowledges funding from DISCOVER-UAH-CM project (Ref.: REACT UE-CM2021-01), 282 cofounded by Community of Madrid (CAM) and European Union (EU), through the European 283 Regional Development Fund (ERDF) and supported as part of the EU’s response to COVID-19 284 pandemic.

## Entry for the Table of Contents

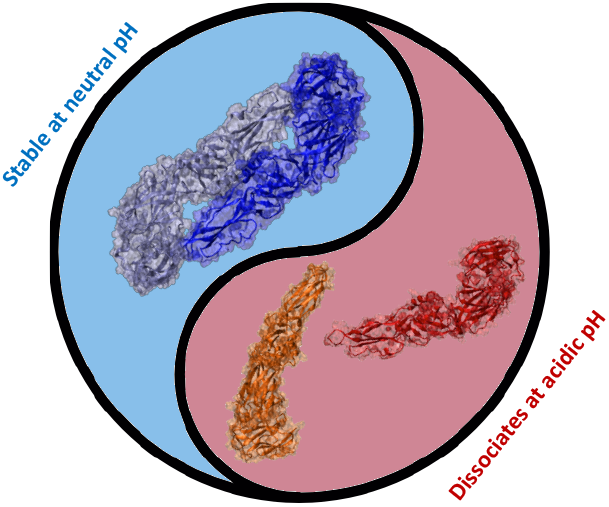

The pH-dependent maturation process in flavivirus is studied with molecular simulations, revealing a crucial role of the sE_2_ complex and its stabilization by the pr protomer in acidic conditions. sE_2_ premature dissociation in the acidic trans Golgi network would hamper the viral maturation, thus pr is a key player in regulating the viral cycle and its destabilization could be regarded as a novel therapeutic strategy against emerging viruses.

Institute and/or researcher Twitter usernames: @AntonioMonari, @elisejdumont, @DrEmmaBignon

