## Supplementary Information for "Molecular basis of the pH-controlled maturation of the tick-borne encephalitis flavivirus"

**Abstract:** Flaviviruses are enveloped viruses causing high public concerns. Their maturation spans several cellular compartments having different pH. Thus, complex control mechanisms are in place to avoid premature maturation. Here we report the dynamical behavior at neutral and acidic pH of the precursor of the membrane fusion protein E of tick-borne encephalitis, showing the different stabilization of the E dimer and the role played by the small fusion-assisting protomer (pr). The comprehension, at atomic resolution, of the fine regulation of viral maturation will be fundamental to the development of efficient strategies against emerging viral threats.

DOI: 10.1002/anie.2021XXXXX

**Table of Contents**


Computational Methods 3

Results and Discussion 3

References 10

### Computational Methods

The initial model of the $(sE/pr)_2$ complexes have been retrieved by the crystal structures obtained by Vaney et al.[1], the missing residues have been recovered by homology model using the Swiss-Model web server.[2] The $sE_2$ dimer was obtained by manually deleting the two pr units from the initial complex. The protonation states of titratable residues were adapted for simulations at acidic pH (5.5) and neutral pH (7.5), according to pKa values as calculated by the propKa3.1 web-server.[3] Potassium counter-ions were added to neutralize each system, which were soaked in a TIP3P water box with a 9.0Å buffer. The ff14SB force field was used, together with the Joung and Cheatham parameters for monovalent ions.[4] Prior to the MD simulation, each system was optimized in three steps coupling 10,000 steps minimization and 12 ns of equilibration at 300K, with decreasing constraints on the protein backbone atoms (scaling factor of 1, 0.5 and 0.1 for the position restraint energy function).

Classical Molecular Dynamics simulations were all performed with the NAMD3 software.[5] For each equilibrated system, a production run of 2 to 2.5 µs was carried out in the NPT ensemble. The Hydrogen mass Repartitioning scheme[6] was used together with RATTLE and SHAKE algorithms [7] in order to use a 4 fs integration timestep. Positions, velocities, energies and pressure values were output every 40 ps. The temperature was kept to 300K and the pressure using the Langevin thermostat with a 1 $ps^{-1}$ collision frequency, and the pressure at 1 atm using the Langevin piston method with an oscillation period of 100 fs and a damping times scale of 50 fs.

Free energy calculations were performed using the MM/GBSA approach as available in the AMBER22 package[8]. The input parameters were kept as such (internal and external dielectric constants of 1 and 78.5, respectively), and the energy decomposition by residue was turned on. In the case of the $(sE)_2$ system in acidic conditions, the binding free energy value was monitored for the three phases of the dimer dissociation, respectively from 0 to 200 ns, from 200 to 1600 ns, and from 1600 ns to the end of the simulation. The $\Delta\Delta G_{binding}$ reported in the main text were computed as the difference between the $\Delta G_{binding}$ in neutral conditions and the one in acidic conditions (last phase of the dissociation for the $(sE)_2$ system).

Figures were rendered using VMD [9] and gnuplot (http://www.gnuplot.info/) softwares.

### Results and Discussion

MM/GBSA calculations were performed in order to estimate the free energy of binding ($\Delta G_{binding}$) between the two sE monomers. The energy decomposition was used in order to pinpoint the residues playing a role in the $(sE)_2$ stability. Results are reported in table **S1** and show that the addition of the pr units stabilizes the $sE_2$ complex in acidic conditions but not in at neutral pH.

The energetic decomposition showing the contribution of each sE amino acids in the total binding energy can be visualized in Figures S1 to S3. The contribution in the $sE_2$ systems with (cpx) and without (apo) the pr units, as well as the difference between the two (delta) allow to highlight the most (de)stabilizating amino acids for the $sE_2$ dimer, in acidic (Figure S1 and S2) and in neutral (Figure S3) conditions.

Interestingly, the change of contribution of the sE residues upon addition of pr is less pronounced in neutral conditions (Figure S3) than with acidic pH (Figure S1). The $sE_2$ binding free energy in acidic condition is much more favorable in the $(pr/sE)_2$ tetramer, with a strong influence of the W101/R316 interaction, while the presence of the pr units limits the destabilizing effect of E7 protonation.

Figure S2 shows the loss of the most important interactions along the MD simulations of $sE_2$ in acidic conditions without the pr units, correlated to the dimer dissociation observed in our simulations.

**Table S1.** Free energy of binding of the two sE monomers with or without pr units, as calculated by MMGBSA. For $(sE)_2$, the values are divided in three phases (0-200ns, 200-1600ns and 1600-end) according to the dynamics of dissociation of the complex. Standard deviation (STD) and standard means of error (SME) are reported. Values are all in kcal/mol.

| System | pH | Simulation time (ns) | $\Delta G_{binding}$ | STD | SME |
|---|---|---|---|---|---|
| $(sE)_2$ | acidic | 0-200 | -49.12 | 14.14 | 1.23 |
| | | 200-1600 | -39.25 | 12.49 | 1.67 |
| | | 1600-end | -18.11 | 15.82 | 2.52 |
| | neutral | all | -83.65 | 14.87 | 1.10 |
| $(sE/pr)_2$ | acidic | all | -44.85 | 9.53 | 1.32 |
| | neutral | all | -67.72 | 12.71 | 1.40 |

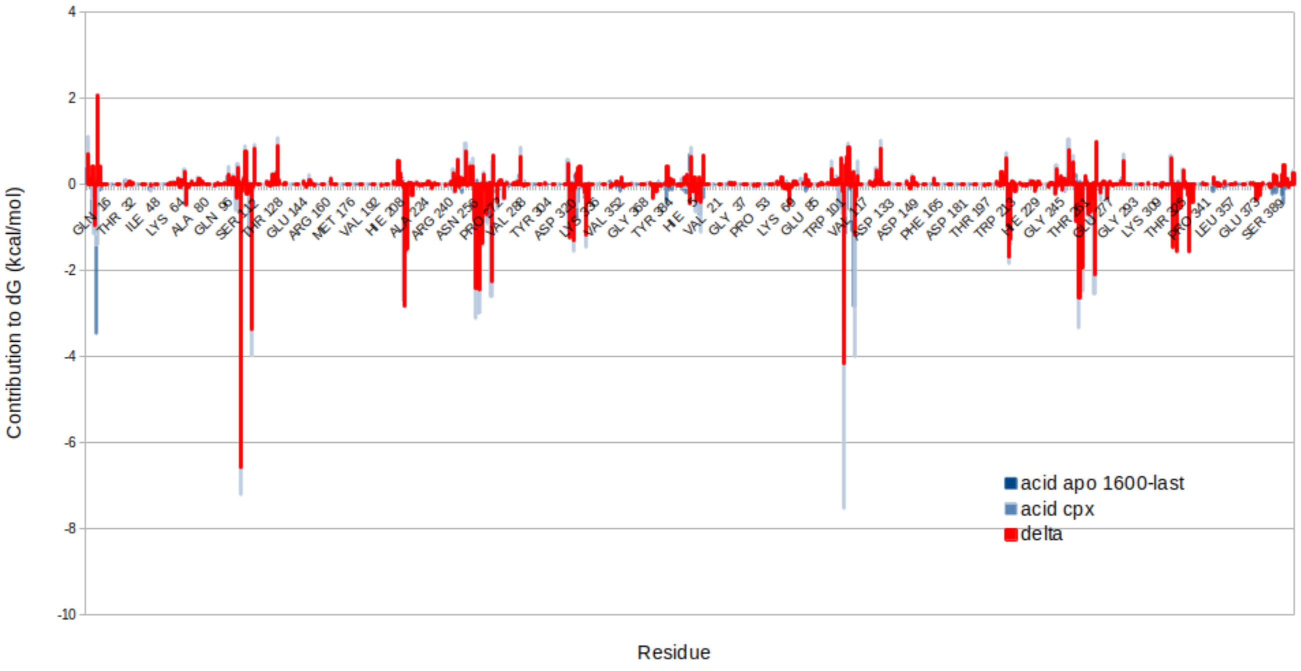


**Figure S1.** Energetic contribution of each sE monomer to the total free energy of binding in acidic conditions, without (dark blue) and with (light blue) the pr units. The difference between both states appears in red in order to highlight the effect of the presence of pr onto the interaction network between sE1 and sE2 residues. One can see that the addition of the pr units tends to enhance stabilizing interactions between the sE monomers.

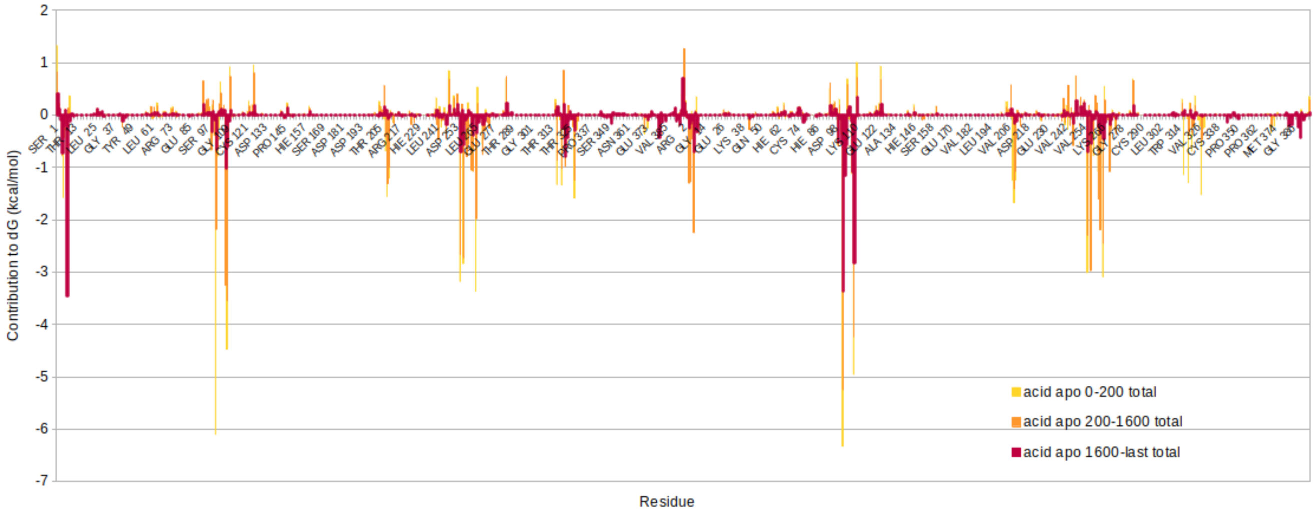


**Figure S2.** Energetic contribution of each sE monomer to the total free energy of binding in acidic conditions without pr units, superimposed for the three phases of the dissociation (0 to 200 ns in yellow, 200 to 1600 ns in orange, 1600 to the end of the simulation in red). One can appreciate the decrease of stabilizing interactions between the two sE monomers as the dissociation takes place.

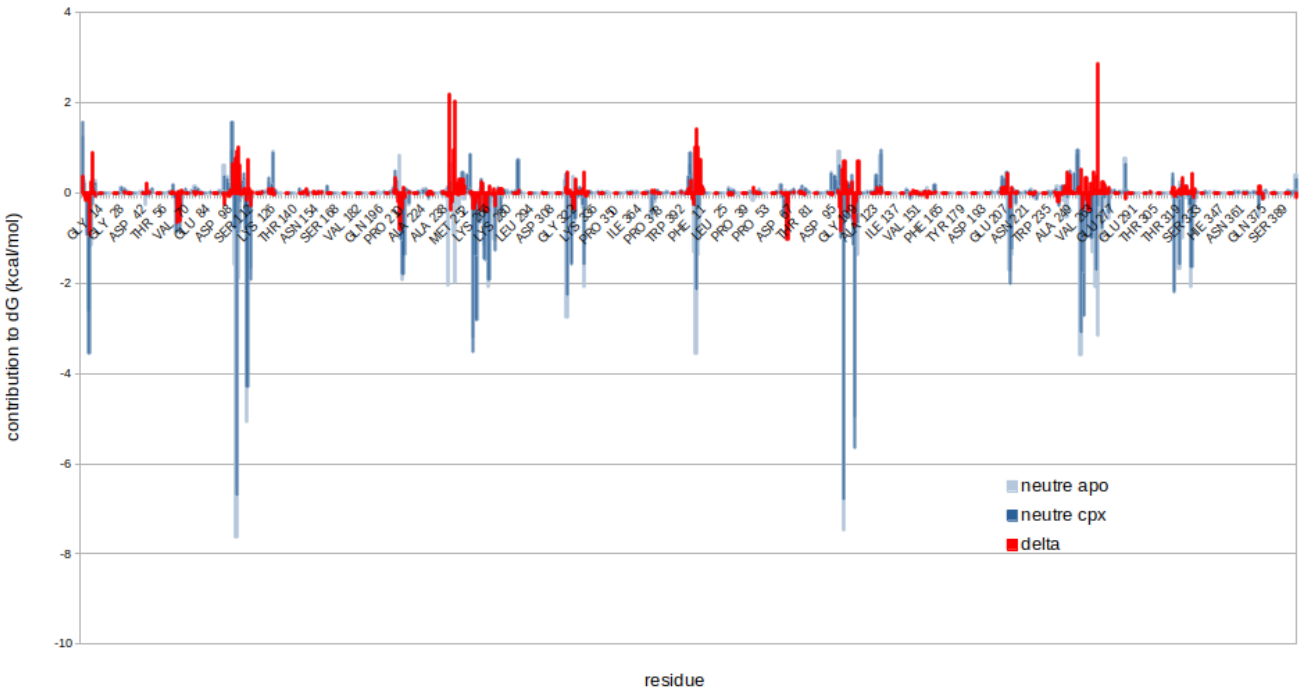


**Figure S3.** Energetic contribution of each sE monomer to the total free energy of binding in neutral conditions, without (dark blue) and with (light blue) the pr units. The difference between both states appears in red in order to highlight the effect of the presence of pr onto the interaction network between sE1 and sE2 residues. One can see that, contrary to what is observed in acidic conditions, the addition of the pr units at neutral pH does not stabilize the $(sE)_2$ dimer.

**Table S2.** Inventory of the most significant non-covalent interactions identified for $sE_2$ and $(sE/pr)_2$ as extracted from the MD simulation. SB strand for salt bridge, HB for hydrogen bond. To measure the inter-residue distance, the following atoms were chosen: for glutamates and aspartates, the carbon atom of the carboxylate group, for lysines, the nitrogens of the ammonium termina is chosen and for arginines, the carbon atom of the guanidium moiety.

| System | Interaction | Nature | Average distance (Å) Acidic pH | Average distance (Å) Neutral pH | Comment |
|---|---|---|---|---|---|
| $(pr/sE)_2$ | K64(sE)-D43(Pr) | SB | 6.15±1.89 (sE1-Pr1);<br>6.24±1.70 (sE2-Pr2) | 8.50±2.02 Å (sE1-Pr1);<br>8.71±2.02 Å (sE2-Pr2) | Transient in MD at acidic pH, disrupted at neutral pH |
| | D67(sE)-S45(Pr) | HB | 4.75±1.56 (sE1-Pr1);<br>6.31±1.78 (sE2-Pr2) | 5.15±1.74 (sE1-Pr1);<br>4.47±1.19 (sE2-Pr2) | Reinforced at acidic pH |
| | H104(sE)-D54(pr) | SB | 5.95±1.18 (sE1-Pr1);<br>5.78±1.02 (sE2-Pr2) | 9.76±2.12 (sE1-Pr1);<br>9.90±2.07 (sE2-Pr2) | Present at acidic pH, mobile, disrupted at neutral pH |
| | D253(sE)-R78(pr) | SB | 5.81±1.77 (sE1-Pr1);<br>4.92±1.03 (sE2-Pr2) | 7.49±1.98 (sE1-Pr1);<br>6.06±1.27 (sE2-Pr2) | Present at acidic pH, mobile, disrupted at neutral pH |
| | A72(sE)-E49(pr) | HB | 5.78±1.02 (sE1-Pr1);<br>5.95±1.18 (sE2-Pr2) | 6.60±0.48 (sE1-Pr1);<br>6.53±0.52 (sE2-Pr2) | Transient |
| | D57(pr)-R316(sE) | SB | 4.57±1.35 (sE2-Pr1);<br>4.95±1.35 (sE1-Pr2) | 3.78±0.72(sE2-Pr1);<br>3.66±0.37(sE1-Pr2) | Reinforced at neutral pH |
| | D57(pr)-S1(sE) | SB | 7.75±2.28 (sE2-Pr1);<br>11.06±3.12 (sE1-Pr2) | 9.31±2.01(sE2-Pr1);<br>8.29±1.23 (sE1-Pr2) | Only transiently, disrupted at neutral pH |
| | D63(pr)-K251(sE) | SB | 3.68±0.31 (sE1-Pr1);<br>3.71±0.36 (sE2-Pr2) | 3.67±0.38 (sE1-Pr1);<br>3.68±0.33 (sE2-Pr2) | Stable |
| | D37(pr)-K251(sE) | SB | 4.95±1.31 (sE1-Pr1);<br>3.90±0.80 (sE2-Pr2) | 6.25±2.17 (sE1-Pr1);<br>4.48±1.39 (sE2-Pr2) | Stable |
| | R160(sE)-E58(pr) | SB | 6.54±1.78 (sE1-Pr2);<br>- disrupted after 20ns<br>8.98±2.62 (sE2-Pr1)<br>- disrupted after 2 μs | 4.30±0.88 (sE1-Pr2);<br>4.48±1.39 (sE2-Pr1) | Disrupted upon protonation of E58 at acidic pH |
| | E7(sE1)-K110(sE2) | SB | 4.03±0.39 (sE2-Pr1);<br>3.95±0.39 (sE1-Pr2) | 4.10±1.16 (sE2-Pr1);<br>4.72±1.19 (sE1-Pr2) | Salt bridge at neutral pH; weakened and disrupted after 2200 ns under acidic conditions |
| | D63(Pr)-K251(sE) | SB | 3.68±0.31 (sE1-Pr1);<br>3.71±0.36 (sE2-Pr2) | | Observed only at acidic pH |
| | D37(Pr)-K251(sE) | SB | 4.95±1.31 (sE1-Pr1);<br>3.90±0.80 (sE2-Pr2) | | Observed only at acidic pH |
| | D8(Pr)-K69(sE) | SB | 8.90±2.62 (sE1-Pr1);<br>10.35±3.00 (sE2-Pr2) | | Observed only at acidic pH |
| $(sE)_2$ | E7(sE)-K110(sE) (*) | SB at neutral pH | 4.03±0.39 Å (sE2-sE1);<br>3.95±0.39 A (sE1-sE2) | 4.10±1.16 Å (sE2-sE1);<br>4.72±1.19 A (sE1-sE2) | |
| | W101(sE)-R316(sE) (‡) | π-cation | 4.49±0.62 Å (sE2-sE1);<br>4.66±0.37 A (sE1-sE2) | 4.04±0.39 Å (sE2-sE1);<br>3.96±0.39 A (sE1-sE2) | |

(*) in absence of protomers, values averaged up to 2.2 μs: 6.63±1.91Å (sE2-sE1) ;8.24±2.60 Å (sE1-sE2) at acidic pH, and 4.78±2.02 Å (sE2-sE1) ;4.40±1.39 Å (sE1-sE2) at neutral pH

(‡) in absence of protomers, values averaged up to 2.2 μs: 10.62±4.97Å (sE2-sE1) ; 6.27±2.41 Å (sE1-sE2) at acidic pH, and 4.48±0.53Å (sE2-sE1) ; 3.98±0.47 Å (sE1-sE2) at neutral pH

During our microsecond range dynamics, two non-covalent interactions for the $sE_2$ dimer are clearly identified: a salt bridge between E7(sE1) – K110(B) – symmetrically E7(B) – K110(A) – and a π-cation interaction between W101(A) and R316(B) -- symmetrically R316(A) – W110(B). The rest of interactions appear minimal, and correspond to hydrophobic, sparse interactions. In particular, no interaction is identified between the two central α-helices shown in Figure S5. These two non-covalent interactions are maintained at neutral pH, yet are disrupted at 2.3 μs at acidic pH, with the protonation of E7. This pH-induced signal leads to an initial weakening of the E7…K110 interchain salt bridge, but also, cooperatively, significantly weakens the π-cation interaction, leading to the opening. Both average distances are significantly higher (by ca. 4 Å) as can be seen from Table S2.

As the protomers become implied in $(sE/pr)_2$, these two interactions exhibit an enhanced stability. The assembly is also strongly, and more importantly, reinforced at sE/Pr interfaces, with a network of three main salt bridges: D37-K251, D63-K251 and R78-D253 that consolidate this network and act cooperatively to consolidate the protein-protein interface. Other non-covalent interactions, including salt bridges but not exclusively, come into play with a more dynamical endeavour.

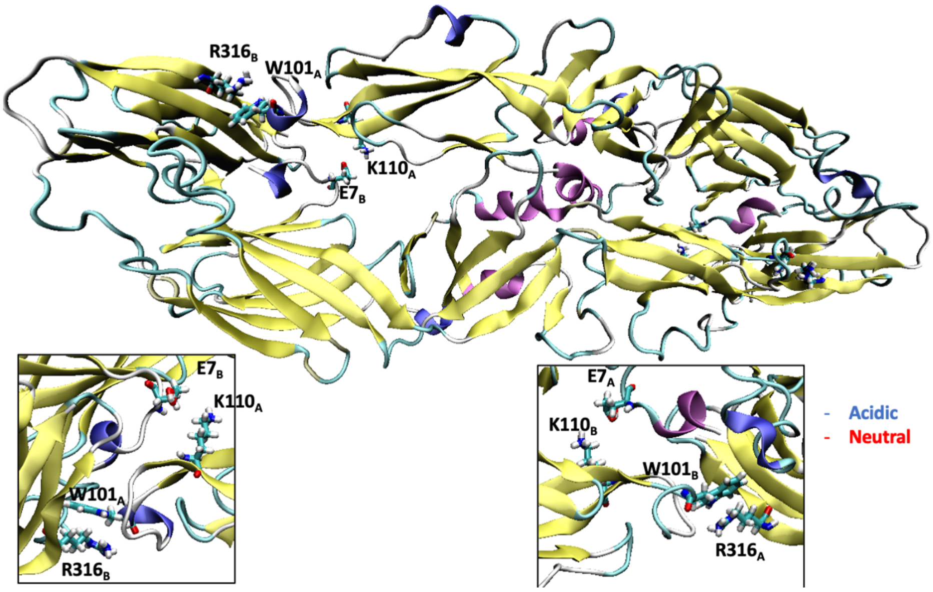


**Figure S4**. Representative snapshot showing the aminoacids responsible for the stabilization of the $sE_2$ interface. The zoom of the two principal interactions are provided as inlays.

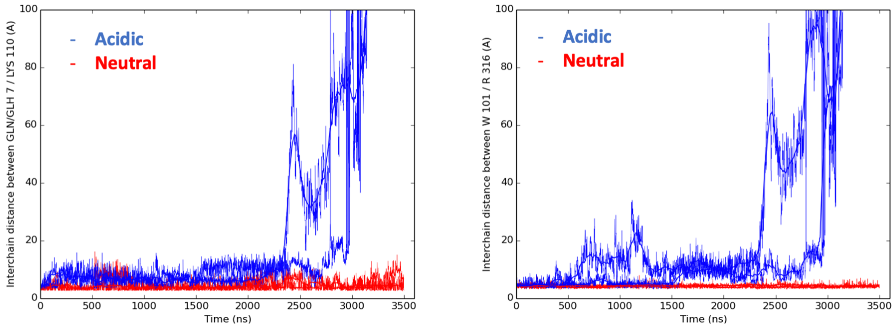


**Figure S5.** Time evolutions of the two non covalent interactions implied in stabilizing the sE2 dimer as shown in Figure S1.

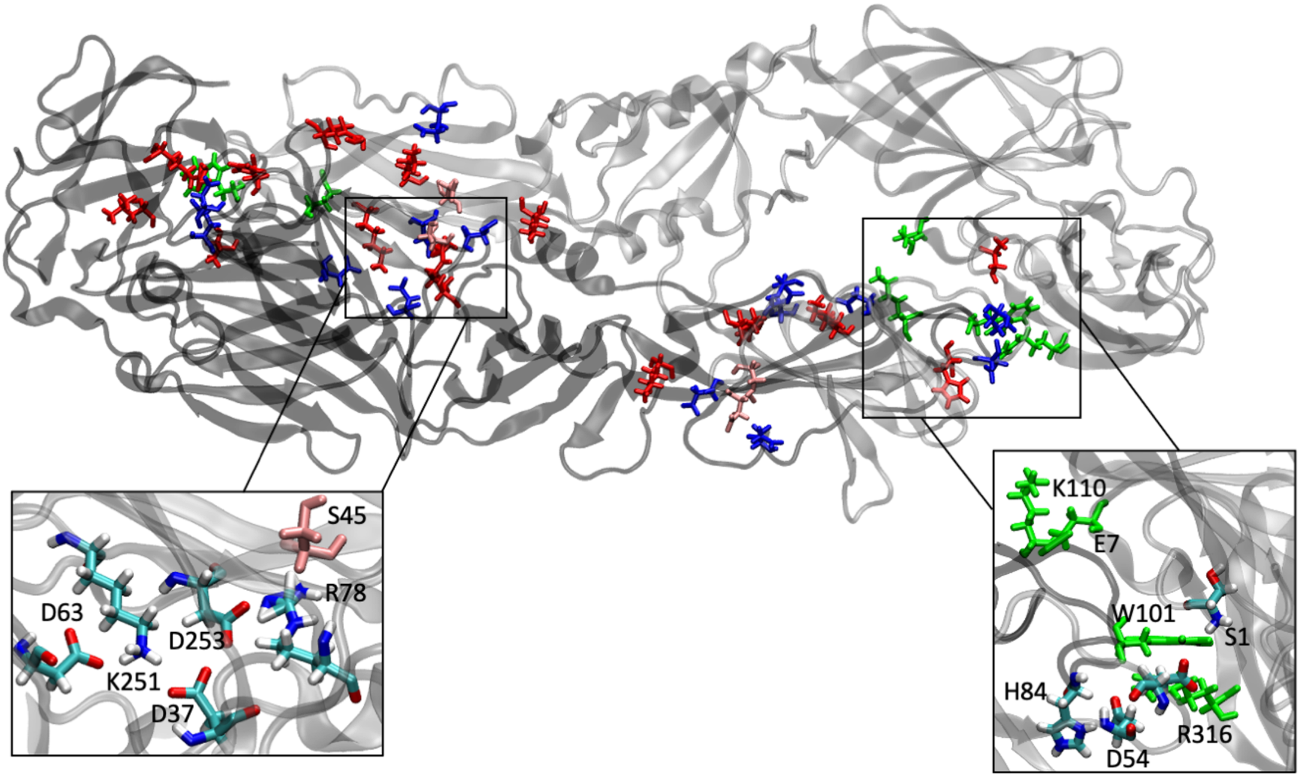


**Figure S5.** Cartoon representation of the (Pr/sE) dimer (Pr1 and sE1 in silver, Pr2 and sE2 in black), with negatively- and positively-charged residues implied in significant salt bridges interactions indicated in blue and ref respectively. A significant non salt-bridge interaction implying S45 and D67, identified on the X-ray structure, is shown in pink. Corresponding inter-residue distances are listed in Table S3.

**Table S3**. Numbering of the pr/sE interactions identified from the crystal structure.

| | Number | Aminoacids |
|---|---|---|
| | 1 | LYS 64 [NZ]/ ASP 43 [OD2] |
| | 2 | ASP 67 [N]/ ASP 43 [O] |
| | 3 | ASP 67 [OD1]/ SER 45 [OG] |
| | 4 | THR 68 [N]/ SER 45 [O] |
| | 5 | THR 68 [OG1]/ SER 45 [O] |
| | 6 | THR 68 [O]/ SER 47 [N] |
| pr1-sE1 'pr2-sE2) | 7 | VAL 70 [N]/ SER 47 [O] |
| | 8 | ALA 72 [N]/ GLU 49 [OE1] |
| | 9 | GLY 102 [O]/ VAL 60 [N] |
| | 10 | ASN 103 [ND2]/ VAL 60 [O] |
| | 11 | HIS 104 [ND1]/ ASP 54 [OD1] |
| | 12 | HIS 104 [ND1]/ THR 52 [O] |
| | 13 | HIS 248 [ND1]/ ASP 61 [OD1] HI |
| | 14 | ALA 249 [N]/ ASP 61 [O] |
| | 15 | ALA 249 [N]/ ASP 61 [OD2] |
| | 16 | LYS 251 [NZ]/ ASP 37 [OD1] |
| | 17 | LYS 251 [NZ]/ ASP 63 [OD2] L |
| | 18 | LYS 251 [NZ]/ TYR 76 [OH] |
| | 19 | ASP 253 [OD1]/ ARG 78 [NH1] |
| | 20 | ASP 253 [OD2]/ ARG 78 [NH1] |
| | 21 | ARG 2 [N] 3.5 GLU 58 [OE1] |
| | 22 | GLU 155 [OE1]/ ARG 67 [NE] |
| | 23 | GLU 155 [OE2]/ ARG 67 [NE] |
| | 24 | GLU 155 [OE2]/ ARG 67 [NH2] |
| | 25 | HIS 157 [ND1]/ GLU 58 [OE1] |
| pr1-sE2 (pr2-sE1) | 26 | HIS 157 [ND1]/ GLU 58 [OE2] |
| | 37 | ARG 160 [NH2]/ GLU 58 [OE1] |
| | 28 | VAL 206 [O]/ ASP 44 [OD2] |
| | 29 | LYS 315 [NZ]/ GLN 55 [O] |
| | 30 | LYS 315 [NZ]/ GLY 56 [O] |
| | 31 | ARG 316 [NH2]/ ASP 54 [OD1] |
| | 32 | ARG 316 [NH1]/ GLU 57 [OE2] |
| | 33 | GLU 329 [OE1]/ GLU 57 [OE2] |
| | 34 | GLU 329 [OE2]/ GLU 57 [OE2] |

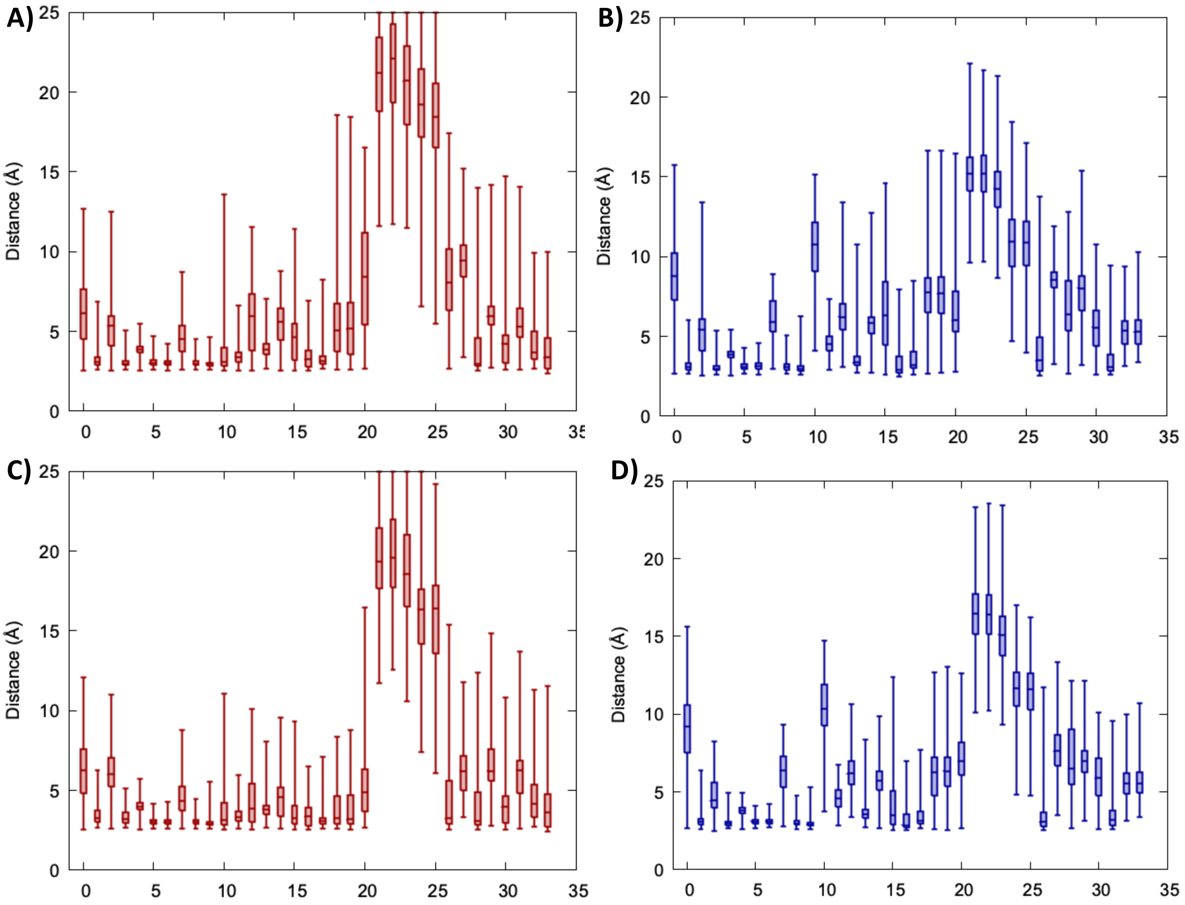


**Figure S6.** Whiskers plot showing the distribution of the distances identified from the crystal structure. For the acidic pr1-sE1 (A) and pr2-sE2 (C) and neutral pr1-sE2 (B) and pr2-sE1 (D).

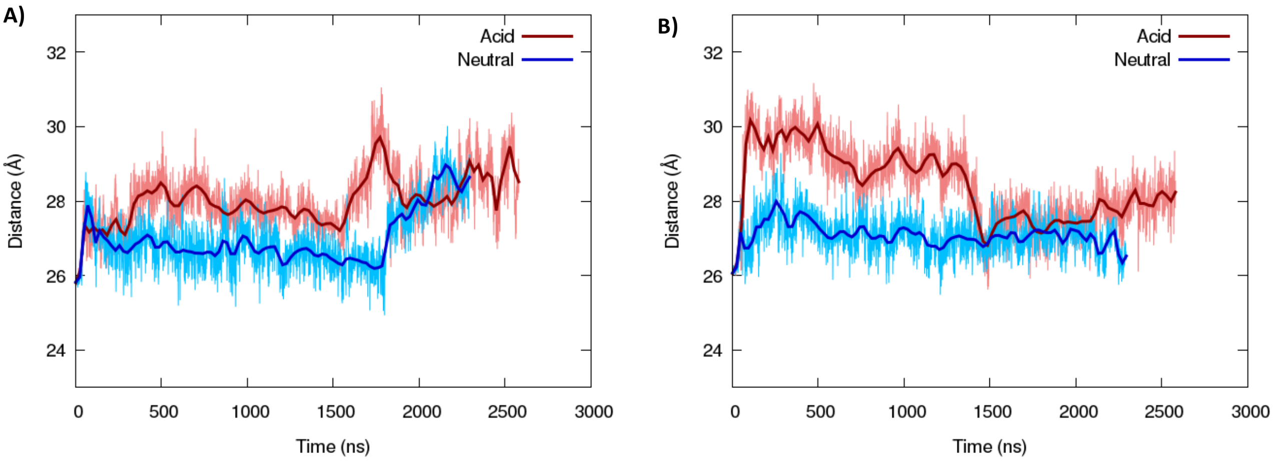


Figure S7. Time series of the distance of the 150-loop from the rest of the sE unit in the $(pr/sE)_2$ tetramer.
